# Clade-specific long-read sequencing increases the accuracy and specificity of the *gyrB* phylogenetic marker gene

**DOI:** 10.1101/2024.07.15.603574

**Authors:** Robert G. Nichols, Emily R. Davenport

## Abstract

Phylogenetic marker gene sequencing is often used as a quick and cost-effective way of evaluating microbial composition within a community. While 16S rRNA gene sequencing (16S) is commonly used for bacteria and archaea, other marker genes are preferable in certain situations, such as when 16S sequences cannot distinguish between taxa within a group. Another situation is when researchers want to study cospeciation of host taxa that diverged much more recently than the slowly evolving 16S rRNA gene. For example, the bacterial gyrase subunit B (*gyrB*) gene has been used to investigate cospeciation between the microbiome and various hominid species. However, to date only primers that generate short-read Illumina MiSeq-length amplicons exist to investigate *gyrB* of the Bacteroidaceae, Bifidobacteriaceae, and Lachnospiraceae families. Here, we update this methodology by creating *gyrB* primers for the Bacteroidaceae, Bifidobacteriaceae, and Lachnospiraceae families for long-read PacBio sequencing and characterize them against established short-read *gyrB* primer sets. We demonstrate both bioinformatically and analytically that these longer amplicons offer more sequence space for greater taxonomic resolution, lower off-target amplification rates, and lower error rates with PacBio CCS sequencing versus established short-read sequencing. The availability of these long-read *gyrB* primers will prove to be integral to the continued analysis of cospeciation between bacterial members of the gut microbiome and recently diverging host species.

## Introduction

Metabarcoding methods have made it possible to study a variety of microbiomes, from environmental to human body sites. These techniques involve sequencing phylogenetic marker genes to characterize the bacteria and archaea present in a microbiome, independent of culturing. The most widely used marker gene for bacteria and archaea is the 16S ribosomal RNA (rRNA) gene [1, 2]. The 16S rRNA gene is an attractive marker sequence because is present in all prokaryotes and archaea [3], includes conserved regions that can be used for primer generation, and multiple hypervariable regions that contain species specific sequences [4]. Sequencing of the 16S rRNA gene can provide very cheap, rapid, and accurate taxonomic identification of the members of a microbiome.

Currently the most popular way to sequence the 16S rRNA gene amplicons is with short-read sequencing technology, like the Illumina MiSeq. Illumina MiSeq sequencing is well validated, very high throughput, and can provide reliable taxonomic information down to the genus level. Unfortunately, 16S Illumina MiSeq sequencing cannot provide species or strain-based resolution [5]. To achieve this level of specificity, one would need to sequence the entire 16S rRNA gene [5]. The 16S gene is only ∼1550 base pairs, which makes it an ideal target for long-read sequencing with PacBio technology [1]. Along with being able to sequence the entire 16S gene, PacBio technology boasts a potential error rate of 1 base pair per million bases after consensus is reached [6]. This error rate is much lower than the reported error rate of 1 out of 1000 bases from Illumina [7]. Having the lowest error rate possible is essential for investigating bacterial strain level differences.

The ability to generate long reads and low error rates is also important for other, non-16S rRNA, marker genes that can be used to investigate other characteristics of a microbiome like evolution and co-evolution. One such target is the bacterial DNA gyrase subunit B gene (*gyrB*), which has been validated and used in studies that require strain and subspecies resolution [8].

This is an attractive marker gene because unlike 16S, there is only one copy of *gyrB* in each bacterial species [9]. Additionally, there are low rates of *gyrB* gene transfer between bacterial species in a microbiome [9] and there is a greater degree of divergence in *gyrB* within and between species when compared other marker genes, like 16S [10]. This makes *gyrB* an ideal marker gene for studies investigating microbiome evolution and co-evolution on relatively short time scales (∼millions of years).

For example, a recent study by Moeller et al. uses bacterial *gyrB* to investigate cospeciation between hominid and three bacterial families within their gut microbiomes: Bacteroidaceae, Bifidobacteriaceae, and Lachnospiraceae [11]. Specifically, the PhyloTAGs software [12] was used to create family-specific primers to amplify the bacterial *gyrB* gene from human, chimp, bonobo and gorilla fecal samples [11]. These amplicons were then sequenced on the Illumina MiSeq and used to create maximum likelihood trees to investigate which bacterial families coevolved with hominids. This short-read data provided a proof of concept that there are potential cospeciation relationships between the bacteria of the gut microbiome and hominid hosts. However, technological limitations in short-read sequencing length and error rates potentially limit the specificity and accuracy of these relationships. To further refine these relationships, updated methodology is needed that will allow for longer, more accurate sequencing of this marker gene.

Sequencing the *gyrB* marker gene with PacBio circular consensus sequencing (CCS) offers two major advantages over short-read sequencing. First, as mentioned above, the ability for PacBio to sequence the full-length gene proves a greater taxonomic resolution when compared to short-read sequencing, which can only sequence a part of the gene [13]. The relatively small gene size of *gyrB* (average 2000-2500 base pairs) makes *gyrB* an ideal target for long-read PacBio sequencing. Second, PacBio CCS typically results in a much lower error rate than what is obtained with short-read sequencing (1 out of a million base pairs vs 1 out of 1000 base pairs) [6, 7]. These lower error rates are necessary for cospeciation studies where single nucleotide variations are used to track bacterial evolution. Studies have already shown that PacBio sequencing of similar sized genes like the 16S gene (1550 base pairs) have error rates close to 0% and produces reads at a single nucleotide resolution [14]. Unfortunately, unlike the 16S gene, there are currently no primers available for long-read *gyrB* sequencing.

In the present study we generated and tested three new *gyrB* primer sets designed for PacBio sequencing for the Bacteroidaceae, Bifidobacteriaceae, and Lachnospiraceae bacterial families. Like the original Moeller study, we used the PhyloTAGs framework to design long-read primers targeted for these three families. We identified primers for each family with the lowest predicted off-target effects. Each primer set was used to generate PacBio sequencing data for several human fecal samples, murine cecal samples, and a mock community. We then compared these results to those obtained with the original short-read primer sets on the same samples. Overall, we were able to see better resolution, less off target amplifications and greater accuracy with the long-read *gyrB* primers when compared to the original short-read *gyrB* primers.

## Materials and Methods

### Creation of long-read *gyrB* primers with PhyloTAGs

To create the new long-read *gyrB* primers for Bacteroidaceae, Bifidobacteriaceae, and Lachnospiraceae families (which will be called the ‘Nichols’ primers going forward), the existing tool PhyloTAGs was used [12]. PhyloTAGs creates primers to amplify a specific gene at a desired level of phylogenetic specificity. A reference database, a 16S rRNA gene (16S) identity map, and a reference organism are needed to use PhyloTAGs. For this project, the reference database includes all the bacterial *gyrB* genes (“gyr_refseq.fa”; n = 31,842 genes in 31,842 bacteria; available on Zenodo, DOI 10.5281/zenodo.10451935) extracted from the NCBI RefSeq non-redundant (nr) database downloaded on April 10, 2023 (“Bacteria.refseq.tar.gz.part1”; available on Zenodo, DOI 10.5281/zenodo.10452184 and “Bacteria.refseq.tar.gz.part2”; DOI 10.5281/zenodo.10452279). Along with the bacterial *gyrB* genes, 16S sequences were extracted from the same species used to create the gyrB database (“16S_refseq.fa”; n = 31,842 genes in 31,842 bacteria; available on Zenodo, DOI 10.5281/zenodo.10451935). A nonredundant version of the corresponding 16S sequences was used to create a 16S rRNA gene identity map with the USEARCH program [15]. This identity map gives a percent identity value for every bacterium used in the database, compared to all other used bacteria. A reference organism is chosen from the bacteria present in the 16S/*gyrB* database based on which bacterial family the primers are being created. For this project, NZ_CP036539 was used as a reference for Bacteroidaceae, NC_008618.1 was used as a reference for Bifidobacteriaceae, and NZ_LR699005 was used as a reference for Lachnospiraceae.

PhyloTAGs requires an ‘idlow’ and an ‘idhigh’ value for primer specificity, to indicate sequence similarity cutoffs for primer generation. For this project, an ‘idhigh’ value of 100 was used for each family. Since sequence similarities of a given taxonomic level differs across the phylogenetic tree [16, 17], we determined ‘idlow’ values for each family separately. The ‘idlow’ values were determined by using PhyloTAGs at various values and testing resulting primers with an *in silico* PCR approach with the USEARCH program [15]. A human microbiome dataset from the NCBI (https://ftp.ncbi.nlm.nih.gov/genomes/HUMAN_MICROBIOM/; downloaded August 8, 2022) was used for a reference database to test the primers. This database includes ∼950 bacterial species specific to the human microbiome, making it an appropriate test database for the primers of interest. Primers with the least amount of off-target amplification were chosen. Ultimately, ‘idlow’ values of 91 for Bacteroidaceae, 95 for Bifidobacteriaceae, and 92 for Lachnospiraceae were selected.

To generate the final candidate primer list, PhyloTAGs was run using the “idhigh” and “idlow” values specified above, a codons value of 7, a consensus value of 90 and the greedy option. The codons value determines the length of the created primers; in this case we are using 7 codons or 21 base pairs in length. The consensus option is the percentage required for a single nucleotide to be used in the primer sequence. Consensus values that fall outside this range get the corresponding IUPAC nucleotide code, e.g. Y is used for a nucleotide that could be C or T. The greedy option is used to explore the entire *gyrB* gene when looking for primers. The primer sequences, degeneracy values and position windows are reported from PhyloTAGs (**Supplemental Tables 1-3**). Ultimately, we chose two primer sets per family based on low degeneracy values (degeneracy values 96 or lower) to test with *in silico* PCR. The best performing primer set (least number of off-target amplifications) was then validated in the lab (**Table 1**).

**Table 1:**
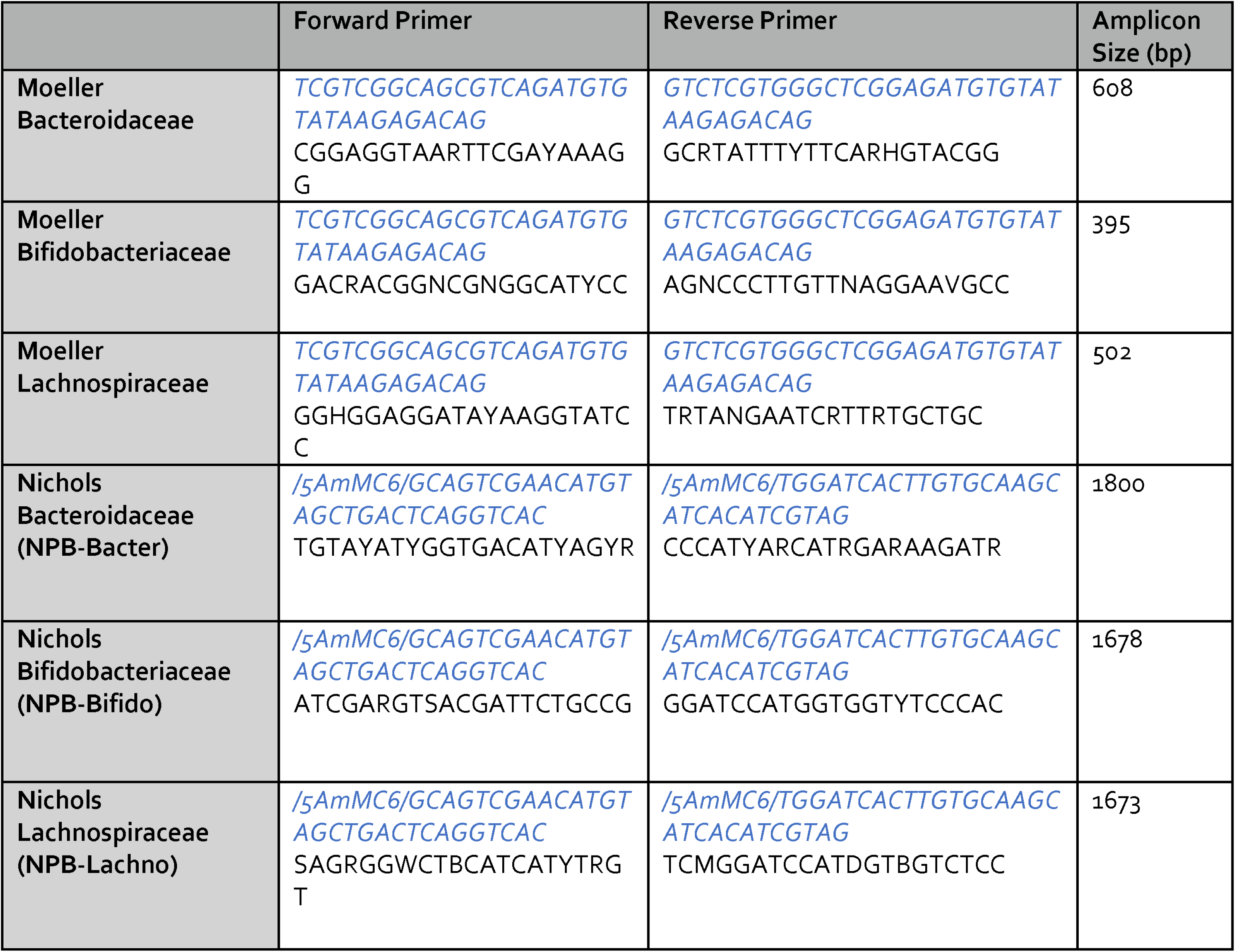
*gyrB* primers and amplicon band sizes. “Moeller” short-read amplicon primer sets were originally described in Moeller et al. [11], while the “Nichols” long-read amplicon primer sets are described here. In the forward and reverse primer columns, blue italicized text shows either the Illumina overhang (Moeller primers) or the PacBio overhang (Nichols primers) needed for sequencing, while black shows the degenerate primers as determined by PhyloTAGs. /5AmMC6/ is a modification needed for PacBio library prep.

### DNA isolation

Five human fecal samples obtained from BioreclamationIVT (now BioIVT), two C57BL/6 wild-type murine cecal samples (generated previously in [18]) and one mock community (ZymoBIOMICS Gut Microbiome Standard, Zymo Research, Irvine, CA) were used to test the efficacy of the primer sets across multiple sample types. DNA was isolated from the 8 test samples using the Omega Bio-Tek E.Z.N.A Stool DNA Kit (Omega Bio-Tek, Norcross, GA) using the manufactures recommended protocol. DNA quantity and quality was measured via a NanoDrop One Spectrophotometer (Thermo Scientific, Waltham, MA).

### *gryB* amplification and sequencing

Detailed protocols for 16S and *gyrB* PacBio library preparation are available on protocols.io (dx.doi.org/10.17504/protocols.io.36wgq31nylk5/v1). Three clade-specific long-read PacBio primer sets (Nichols, generated here) and three short-read primer sets (Moeller, previously published [11]) were used to amplify *gyrB* of Bacteroidaceae, Bifidobacteriaceae, and Lachnospiraceae as these were the three families previously investigated by Moeller et al. (**Table 1**). DNA from the five human samples (H1-H5), two murine samples (M1 and M2) and mock sample (GM) described above underwent library preparation for *gyrB* sequencing. PCR reactions contained Platinum SuperFi DNA polymerase master mix (Invitrogen), 0.4 M of both forward and reverse primers, and 10 ng of template DNA for each sample. PCR conditions for the PacBio *gyrB* amplifications were: 30 sec at 95 °C; 30 cycles of 30 sec at 95 °C, 30 sec at 57 °C and 1 min at 72 °C; then 5 min at 72 °C. PCR conditions for the original Moeller MiSeq *gyrB* amplifications were: 2 mins at 98 °C; 25 cycles of 10 sec 98 °C, 20 sec 56.6 °C, 15 sec 72 °C; then 1 cycle of 5 min at 72 °C, as previously described [19]. PCR reactions were cleaned up using SPRI magnetic beads (Clean NGS) as described in the manufactures provided instructions. Samples were then run on a 1x agarose gel to visualize correct amplification size, between 1600-1800 bp for PacBio amplicons and between 300-700 bp for MiSeq amplicons (**Table 1**). DNA was transferred to the Pennsylvania State Genomics Core facility where amplicon size selection was performed with a BluePippin (Sage Science, Beverly, MA), indexes were added on to each sample, and then samples were mixed in equimolar amounts using SequalPrep Normalization Plates (ThermoFisher Scientific, Waltham MA). The libraries created with the original Moeller primers were sequenced on the Illumina MiSeq with 250x250 paired-end sequencing using Reagent Kit v2, resulting in ∼310,000 reads per sample. The amplicons created with the new PacBio primers were sequenced on the PacBio Sequel lle using the SMRTbell Express Template Prep Kit 2.0 and the Sequel II Binding Kit, resulting in ∼100,000 reads per sample. The library prep schemes for both PacBio and Illumina MiSeq amplicons can be seen in **Figure 1**.

**Figure 1:**
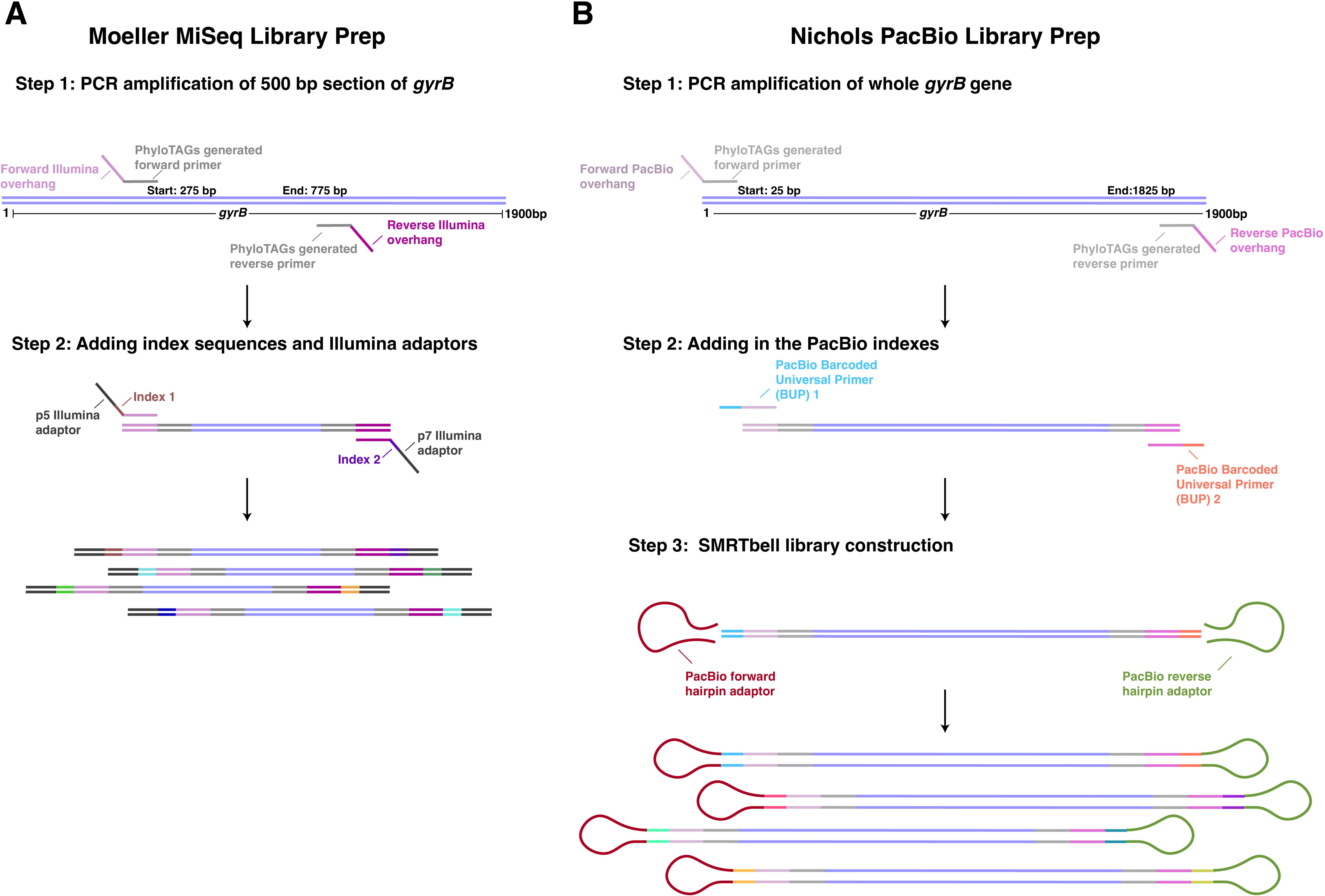
Library preparation schemes for MiSeq and PacBio. A) Library preparation scheme for MiSeq sequencing of *gyrB* amplicons using the Moeller short-read primers. Briefly, family specific primers (gray) with Illumina overhang sequences (pink/purple) are used to amplify a ∼500bp amplicon from the *gyrB* gene. In a second step, Illumina p5/p7 adapters (dark gray) and index sequences (multi-colored) are added to each sample. B) Library preparation scheme for the PacBio sequencing of the *gyrB* amplicons using the Nichols long-read primers. Briefly, family specific primers (gray) with PacBio adapter sequences (light/dark pink) are used to amplify an ∼1800bp amplicon from the *gyrB* gene. In Step 2, the PacBio barcoded universal primers are added to either end (pink/blue and dark pink/salmon) of each amplicon. Finally, in Step 3, the PacBio hairpin adapters (maroon/green) are added to circularize the fragments for circular consensus sequencing (CCS). Ultimately, each library preparation scheme results in amplicons indexed by sample that can be multiplexed for sequencing.

### 16S amplification and sequencing

16S sequencing of the five human samples (H1-H5), two murine samples (M1 and M2) and mock sample (GM) was also performed on both the Illumina MiSeq and PacBio Sequel IIe. PCR reactions contained Platinum SuperFi DNA polymerase master mix (Invitrogen), 0.4 M of both forward and reverse primers, and 10 ng of template DNA for each sample. Traditional long-read 16S primers (27F: AGRGTTYGATYMTGGCTCA and 1492R RGYTACCTTGTTACGACTT) were used for PacBio sequencing [20]. PCR conditions for the PacBio 16S amplifications were: 30 sec at 95 °C; 30 cycles of 30 sec at 95 °C, 30 sec at 57 °C and 1 min at 72 °C; then 5 min at 72 °C. For MiSeq sequencing, V4 primers were used for 16S sequencing (515F: GTGYCAGCMGCCGCGGTAA and 806R: GGACTACNVGGGTWTCTAAT) [21]. PCR conditions for the MiSeq 16S samples were: 2 mins at 98 °C; 25 cycles of 10 sec 98 °C, 20 sec 56.6 °C, 15 sec 72 °C; then 1 cycle of 5 min at 72 °C [19]. PCR reactions were cleaned up using SPRI magnetic beads (Clean NGS) as described in the manufactures provided instructions. Samples were then run on a 1x agarose gel to visualize correct amplification size, ∼1400 bp for PacBio 16S amplicons and ∼350 bp for MiSeq 16S amplicons. DNA was transferred to the Pennsylvania State Genomics Core facility where amplicon size selection was performed with a BluePippin (Sage Science, Beverly, MA), indexes were added on to each sample, and then samples were mixed in equimolar amounts using SequalPrep Normalization Plates (ThermoFisher Scientific, Waltham MA). The libraries created with the V4 primers were sequenced on the Illumina MiSeq with 250x250 paired-end sequencing using Reagent Kit v2 with the *gyrB* MiSeq samples (described above), resulting in ∼310,000 average reads per sample. The long-read 16S amplicons were sequenced on the PacBio Sequel lle with the *gyrB* PacBio samples (described above) using the SMRTbell Express Template Prep Kit 2.0 and the Sequel II Binding Kit, resulting in ∼100,000 average reads per sample.

### Analysis of *gyrB* sequencing reads with DADA2

Both the Illumina and PacBio *gyrB* reads were analyzed with DADA2 (v 1.26) [22]. Briefly, for the PacBio data, we used minQ=3, minLen=1000, maxLen=2000, maxN=0, rm.phix = FALSE, and maxEE=3 for the filter and trim step. Samples were then dereplicated and denoised with an error rate created from the samples. Chimeras were then removed with the consensus method and samples were classified with the custom *gyrB* reference database that used in primer creation (“NCBI_nr_gyrB”). Unclassified reads were assigned with BLASTX and the RefSeq NR database to obtain taxonomy. The top hit for each read with a percent identity above 75% was used for taxonomy assignment. BLASTX results with a percent identity of less than 75%, or a non-*gyrB* gene assignment resulted in an ‘Unclassified’ taxonomy assignment.

For the MiSeq data, reads were filtered with truncLen=c(250,230), maxN=0, maxEE=c(2,2), truncQ=2, rm.phix=TRUE, and trimLeft being variable, depending on the size of the primers, see *dada2_gyrB_analysis.Rmd* in the GitHub repository. Samples were used to create an error rate and then denoised and merged as described above. Chimeras were removed from the merged reads. Like the PacBio reads, the MiSeq reads were classified with the custom *gyrB* reference database (“NCBI_nr_gyrB”). Unclassified reads were assigned with BLASTX, with results at a percent identity of less than 75% or a non-GyrB protein assignment resulted in an ‘Unclassified’ taxonomy assignment. In addition, MiSeq reads were also classified with the corresponding PacBio reads as a database to check for overlap.

### Creation of pseudo-MiSeq sequences and comparison to PacBio ASVs

PacBio *gyrB* reads were cut using the Moeller short-reads primers and the *in silico* PCR option with the USEARCH program [15]. The resulting cut reads were treated as “pseudo-MiSeq” amplicons from the same samples, to mimic the specificity that a shorter amplicon would provide. These cut reads were analyzed the same way as the original PacBio reads described above. In addition to classifying these reads with the custom *gyrB* reference database (“NCBI_nr_gyrB”), the pseudo-MiSeq reads were also classified with the original parent PacBio ASVs. This second classification was done to match the pseudo-Miseq read with its parent PacBio ASV read. In some cases, the pseudo-Miseq read matched to multiple parent PacBio ASVs, this resulted in a ‘NA’ classification. These reads were individually searched and matched based on similar read count data between the parent PacBio ASV and the pseudo-Miseq ASV. Additionally, some low abundance (<0.1% total abundance) pseudo-MiSeq reads did not have a parent PacBio ASV match, these were denoted as ‘unassigned’.

The pseudo-MiSeq ASV sequences and their matching Pacbio ASV sequences were then individually aligned using the ClustalW algorithm within the MEGA 11 program [23]. After alignment, maximum likelihood trees were created with MEGA 11 using the general time reversible (GTR) model, including invariant sites and used the nearest neighbor interchange heuristic method [23]. The resulting trees were then imported to dendroscope to create tanglegrams to visualize matches of pseudo-MiSeq sequences with their corresponding parent PacBio sequence [24]. The environment for tree exploration (ETE) toolkit was used to calculate the normalized Robinson-Foulds (RF) symmetric difference between each tree pair (parent PacBio ASVs and pseudo-MiSeq ASVs) [25].

## Results

To generate the ideal long-read sequencing amplicons of *gyrB* for phylogenetic comparisons, we first designed multiple sets of primers per bacterial family considered. We evaluated those primer pairs *in silico* for off-targets effects, and then the best performing pairs were validated experimentally using human and murine cecal samples as well as a mock community of known composition (ZymoBIOMICS Gut Microbiome Standard). These results were then compared to data generated on the MiSeq using the Moeller short-read primers. For each bacterial family examined (Bacteroidaceae, Bifidobacteriaceae, and Lachnospiraceae), we first identified the top performing long-read primer pair. We then showed that the long-read primers demonstrated better phylogenetic resolution when compared to the corresponding short-read primer pair for that family.

### PhyloTAGs primer locations are chosen based on low degeneracy locations

We ran PhyloTAGs on each bacterial family to generate a list of candidate long-read primers (see Methods). The results from PhyloTAGs were collapsed based on the degeneracy score to show the locations with the lowest degeneracy values (**Figure 2).** Nichols long-read primers were chosen in spots of low degeneracy to give the longest possible read with the fewest potential predicted off-target amplifications (**Table 1**). For example, the Bifidobacteriaceae degeneracy plot has a low degeneracy primer region that could result in a longer read (**Figure 2B**), but after *in silico* PCR, it was determined that the indicated primer locations resulted in the fewest off-target amplifications.

**Figure 2:**
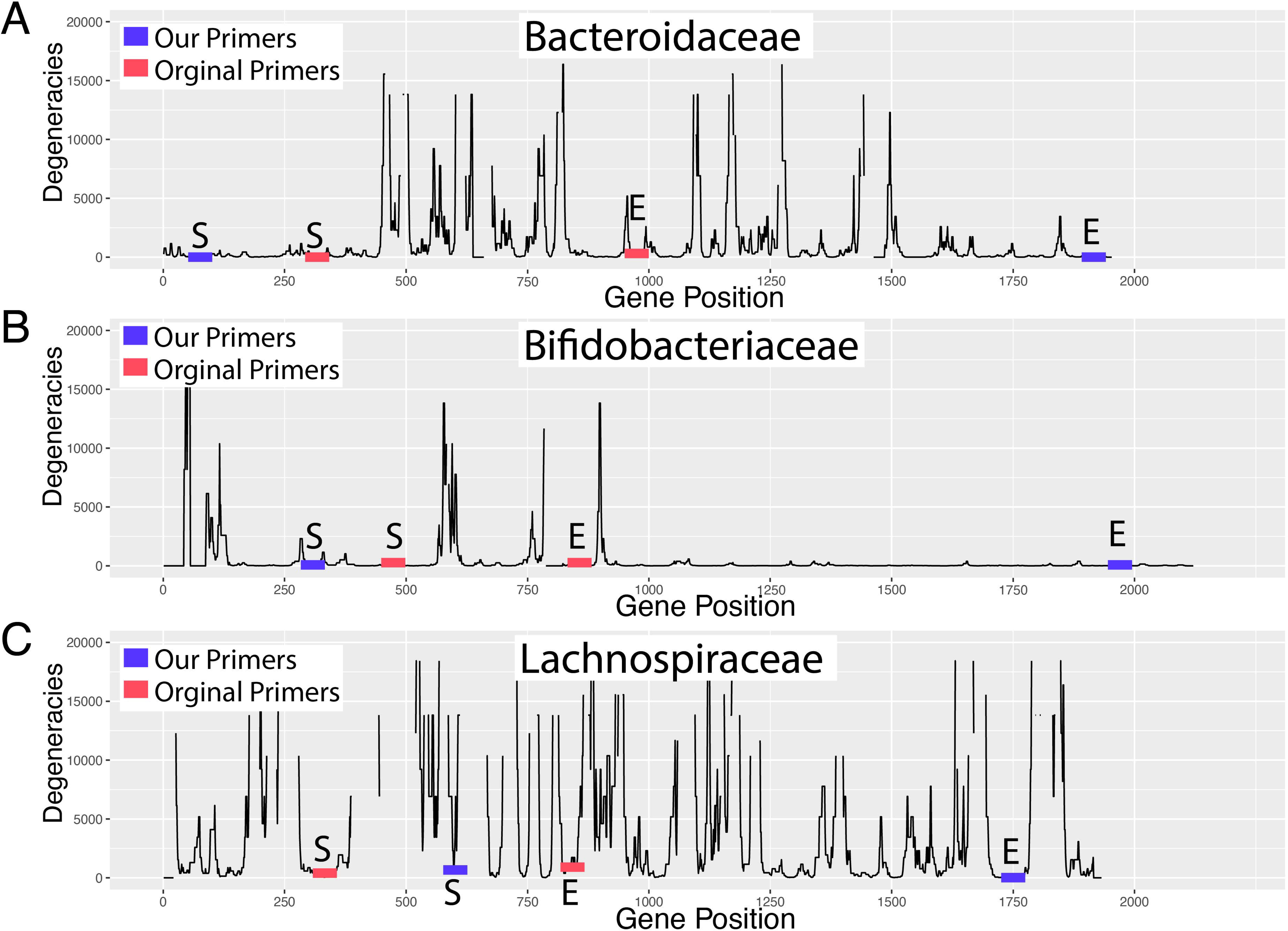
Degeneracy plots showing the genetic coordinates of the Nichols long-read and Moeller short-read *gyrB* primers. We developed long-read, clade specific primers for the bacterial families A) Bacteroidaceae, B) Bifidobacteriaceae, and C) Lachnospiraceae for use with PacBio sequencing (blue). They target a longer region of the *gyrB* gene (x-axis) in each family to capture more of the degeneracies between strains (y-axis) than primers designed for short-read sequencing (red). Number of degeneracies per site is indicated on the y-axis for each family. S = start, E = end.

It should be noted that *gyrB* gene length varies both between and within species. Within our complete assembled database which spans the full bacterial phylogeny, lengths range from 1352 base pairs (bp) to 3246 bp, with the average size ranging from 2000 bp to 2500 bp (**Figure 3**). Bacterial *gyrB* also varies in length within each of the three bacterial families utilized for this study. In Bacteroidaceae (n = 39 representative species), *gyrB* size ranges from 1959 bp to 1974 bp, with the average size being between 1959 bp and 1962 bp. Bifidobacteriaceae had the most number of representative species in the *gyrB* database (n = 146) and had *gyrB* sizes ranging from 2040 bp to 2172 bp and an average length of 2091bp. Lachnospiraceae had the least number of representative species in the *gyrB* database (n = 6) and had *gyrB* lengths that varied between 1911 bp and 2244 bp, with an average size between 1911 bp and 1938 bp.

**Figure 3:**
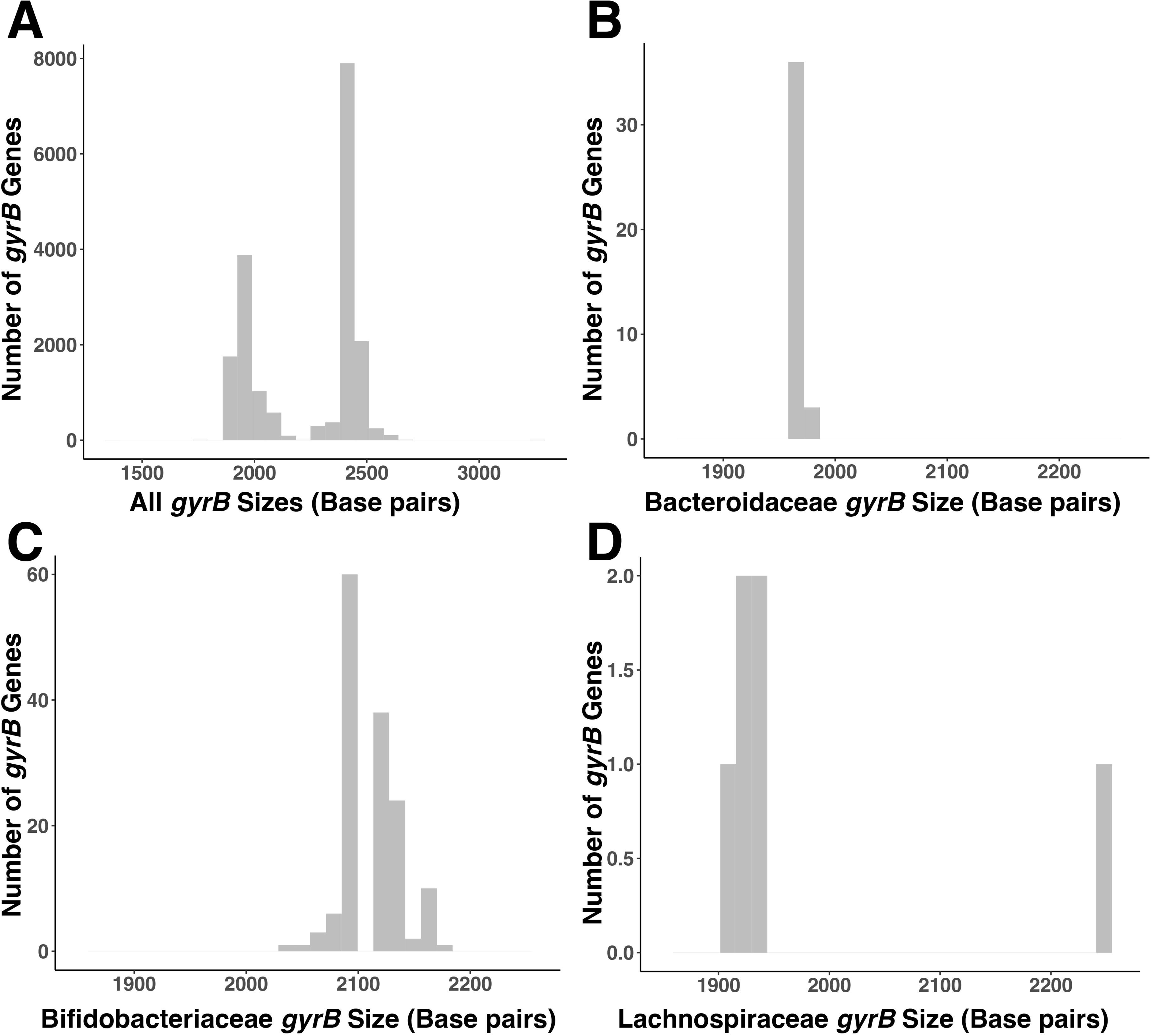
Range of *gyrB* gene sizes within the *gyrB* reference database per family. The length of the *gyrB* gene varies across taxa included in the *gyrB* reference database A), including the Bacteroidaceae B), Bifidobacteriaceae C), and Lachnospiraceae D) families.

### PacBio *gyrB* ASVs result in higher specificity and accuracy than the pseudo-MiSeq *gyrB* ASVs

Theoretically, the longer sequencing lengths and higher fidelity of PacBio reads (created from the Nichols long-read primers) compared to shorter Illumina reads (created from the Moeller short-read primers) should result in both more sequence space and more accurate variant calls for phylogenetic tree construction. To demonstrate this is the case, we amplified and sequenced *gyrB* amplicons from a mock community (GM), human fecal samples (H1-H5), and murine samples (M1 and M2) using both the Nichols long-read and Moeller short-read primer pairs (see Methods).

First, to establish the improved performance of the high fidelity long-reads, we compared the amplicons generated via PacBio sequencing to what we call “pseudo-MiSeq” reads. The pseudo-MiSeq reads were generated by trimming each raw PacBio read to its corresponding MiSeq read by using *in silico* PCR and the Moeller short-read primers [11]. The resulting tanglegram for the Bacteroidaceae family (**Figure 4**) shows good concordance between the long PacBio and pseudo-MiSeq short read, with 28 out of 38 pseudo-MiSeq ASVs classifying to the same bacterial species as their parent PacBio ASV and a normalized Robinson-Foulds distance (nRF) of 0.39. Of the 10 pseudo-MiSeq ASVs that do not match to their parent PacBio ASVs, 6 (or 15.7% of the total ASVs) are unassigned, 3 (or 7.9% of the total ASVs) classify to a different bacterial species than their parent PacBio ASVs and 1 (or 2.6% of the total ASVs) is classified at a less specific taxonomic level than their parent PacBio ASVs (**Figure 4**).

**Figure 4:**
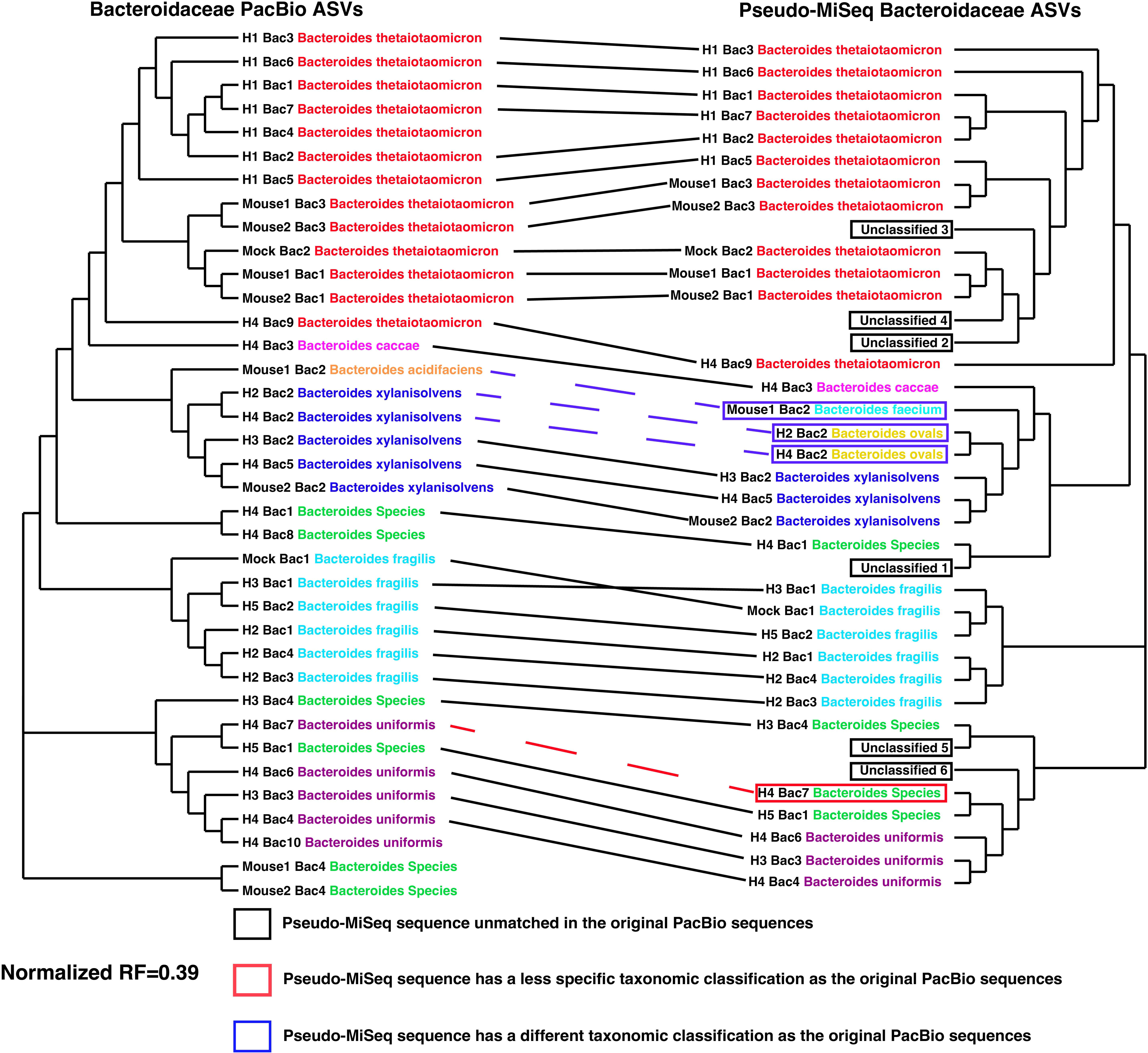
Tanglegram of *gyrB* maximum likelihood trees comparing the Bacteroidaceae PacBio ASVs and the ‘pseudo-MiSeq’ ASVs. Bacteroidaceae phylogeny was generated using the maximum likelihood (ML) method on ASV data collected across all samples. ML trees for long-read PacBio ASVs (Left) and short-read pseudo-MiSeq ASVs (Right) were compared with a tanglegram. Each tip is annotated with the sample it originated from, the ASV number, and the taxonomic information after classification. ASV classification names are colored by species. The solid black lines connecting the two trees illustrate a 1 to 1 relationship where the pseudo-MiSeq ASV results in the same classification as the PacBio parent read. The red dashed line illustrates a scenario where the pseudo-MiSeq sequence has a less specific classification than the parent PacBio sequence. The blue dashed line represents conflicting classifications between the pseudo-MiSeq ASV and its parent PacBio ASV. The normalized Robinson-Fould distance for these two Bacteroidaceae trees was 0.39.

There is also somewhat decent concordance between the long PacBio and pseudo-MiSeq short reads for Bifidobacteriaceae, with 10 out of 22 pseudo-MiSeq ASVs classifying to the same bacterial species as their parent PacBio ASV and a nRF of 0.33 (**Supplemental Figure 1)**. Of the 12 ASVs that do not match their parent PacBio ASVs, 5 (or 22.7% of the total ASVs) are unassigned, 5 (or 22.7% of the total ASVs) classify to a different bacterial species than their parent PacBio ASVs, and 2 (or 9% of the total ASVs) are classified at a less specific taxonomic level than their parent PacBio ASV.

Lachnospiraceae shows the least concordance between the long and pseudo-MiSeq short reads, with 36 out of 59 pseudo-MiSeq ASVs classifying to the same bacterial species as their parent PacBio ASV and a nRF of 0.40 (**Supplemental Figure 2)**. Of the 23 ASVs that do not match their parent PacBio ASVs, 9 (or 15.2% of the total ASVs) are unassigned, 12 (or 20.3% of the total ASVs) classify to a different bacterial species than their parent PacBio ASVs and 2 (or 3.3% of the total ASVs) are classified at a less specific taxonomic level than their parent PacBio ASV.

Overall, the amplicons generated from Nichols long-read primers resulted in more specific taxonomic assignments when compared to the pseudo-MiSeq sequences. In all three bacterial families tested, we saw unassigned ASVs, incorrect taxonomic assignment of ASVs, and less specific taxonomic assignments for the pseudo-MiSeq ASVs when compared to their parent PacBio amplicons. These discrepancies were most visible in the Lachnospiraceae family which had a nRF=0.40, followed by Bacteroidaceae with a nRF=0.39, and then Bifidobacteriaceae with a nRF of 0.33.

### Improved performance of accurate, long-read *gyrB* amplicons vs. short-read *gyrB* amplicons

The pseudo-MiSeq read analysis above demonstrates that the added amplicon length obtained with the PacBio reads provides useful phylogenetic information compared to short reads of equal quality. To further assess the breadth and specificity of the amplicons, as well as the effect of sequencing accuracy, we amplified the mock (GM), human (H1-H5), and murine (M1 and M2) samples described above using the Moeller short-read primers and sequenced using Illumina MiSeq technology [11]. We first assessed the extent of off-target amplification using each primer set **(Table 2).** The Moeller short-read primers for Bacteroidaceae resulted in 82 ASVs across all eight samples. Of the 82 ASVs, 55 ASVs (98.91% of total reads) classified to Bacteroidaceae (**Table 2, Supplemental Figure 3, Supplemental Table 4)**, for an off-target rate of 1.09%. The Nichols long-read Bacteroidaceae primers resulted in 31 ASVs (100% of total reads) across the same samples with all 31 ASVs classifying to Bacteroidaceae and a total off-target rate of 0% (**Table 2, Supplemental Figure 4, Supplemental Table 4**). Therefore, the proportion of usable, on-target data was much higher with the new long-read primers (100%) vs. the established short-read primers (98.91 %) for Bacteroidaceae.

**Table 2:** Performance of each primer set across samples. “On target” is defined as reads/ASVs classifying to the bacterial family targeted by the primer. “Off target” is defined as reads/ASVs classifying to any other bacterial family. A breakdown of classified ASVs and total reads are reported summed across all samples tested. Additionally, an off-target rate is the percent of off target reads for each primer set. The Nichols long-read primers consistently show lower off target amplification rates than the Moeller short-read primers.

| Primer set | Total ASVs | On Target ASVs | Off Target ASVs | Total Reads | On Target Reads | Off Target Reads | Off Target Rate |
| --- | --- | --- | --- | --- | --- | --- | --- |
| Moeller Bacteroidaceae | 82 | 55 | 27 | 812985 | 804113 | 8872 | 1.09% |
| Nichols Bacteroidaceae | 31 | 31 | 0 | 346547 | 346547 | 0 | 0.00% |
| Moeller Bifidobacteriaceae | 256 | 28 | 228 | 707401 | 610610 | 96791 | 13.68% |
| Nichols Bifidobacteriaceae | 66 | 26 | 40 | 262096 | 259754 | 2342 | 0.89% |
| Moeller Lachnospiraceae | 367 | 250 | 117 | 1170397 | 898401 | 271996 | 23.24% |
| Nichols Lachnospiraceae | 456 | 326 | 130 | 693009 | 620713 | 72296 | 10.43% |

Using only the forward Moeller Bifidobacteriaceae reads for classification (as was done in Moeller et al. [11]) resulted in 256 ASVs across the five samples that showed amplification (ZymoBIOMICS Gut Microbiome Standard [GM], Human Fecal Sample 1 [H1], Human Fecal Sample 2 [H2], Human Fecal Sample 4 [H4], and Human Fecal Sample 5 [H5]). Of those 256 ASVs, only 28 ASVs (representing 86.32% of total reads) were classified to Bifidobacteriaceae (**Figure 5**, **Table 2, Supplemental Table 5**). This resulted in a total off-target rate of 13.68%. The Nichols long-read Bifidobacteriaceae primers resulted in 66 total ASVs across the same samples, with 26 ASVs (representing 99.11% of total reads) classifying to Bifidobacteriaceae and a total off-target rate of 0.89% (**Table 2, Supplemental Figure 5, Supplemental Table 5**). Therefore, the proportion of usable, on-target data was much higher with the Nichols long-read primers (99.11%) vs. the Moeller short-read primers (86.32%) for Bifidobacteriaceae.

**Figure 5:**
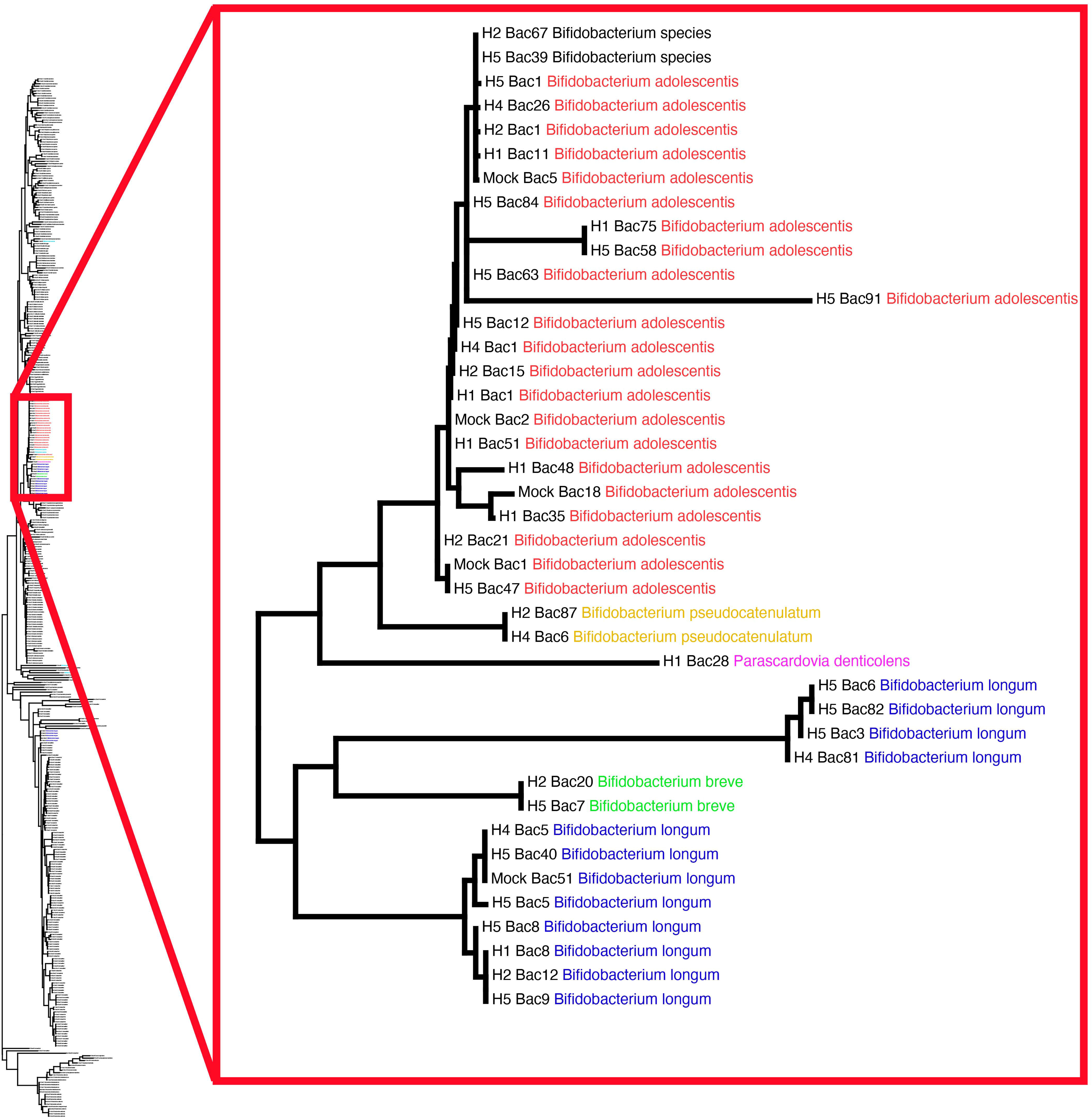
The *gyrB* maximum likelihood tree for Bifidobacteriaceae created from short-read MiSeq ASVs demonstrates extensive off-target amplicons. 256 *gyrB* ASVs were classified from the Bifidobacteriaceae MiSeq reads across the eight samples sequenced. Tips of the tree are ASVs from each sample, labeled first with the sample identifier they came from, the ASV label, and the taxonomic classification of that ASV. ASVs from the same Bifidobacterium species are colored identically for visualization, with off target ASVs colored in black. The complete ML tree containing all 256 ASVs can be seen on the left. Of the 256 *gyrB* ASVs, only 28 ASVs classified to Bifidobacteriaceae. The majority of the ASVs that classified to Bifidobacteriaceae can be seen in the punch-out on the right.

The Moeller short-read primers for Lachnospiraceae resulted in 367 ASVs across the eight samples. Of the 367 ASVs, only 250 ASVs (76.76 % of total reads) were classified to Lachnospiraceae (**Table 2, Supplemental Figure 6, Supplemental Table 6)**. This resulted in a total off-target rate of 23.24%. The Nichols long-read Lachnospiraceae primers resulted in 456 ASVs with 326 ASVs (89.57%) classified to Lachnospiraceae and a total of target rate of 10.43% (**Table 2, Supplemental Figure 7, Supplemental Table 6)**. Therefore, the proportion of usable, on-target data was much higher with the new long-read primers (89.57%) vs. the established short-read primers (76.76 %) for Lachnospiraceae.

Overall, the Nichols long-read primers result in a lower off-target amplification rate when compared to the Moeller short-read primers. The Nichols Bifidobacteriaceae long-read primers in particular resulted in 99.1% classified ASVs, compared to the 86.3% classification rate with the amplicons created from the Moeller short-read primers, for an increase of 12.8% accuracy.

### 16S data confirms presence of each bacterial family in each sample

16S data was used to confirm the presence or absence of Bacteroidaceae, Bifidobacteriaceae, and Lachnospiraceae in the samples used. We detect ASVs for Bacteroidaceae and Lachnospiraceae in all samples (**Supplemental Table 7**). We only see Bifidobacteriaceae ASVs in samples ZymoBIOMICS Gut Microbiome Standard (GM), Human Fecal Sample 1 (H1), Human Fecal Sample 2 (H2), Human Fecal Sample 4 (H4) and Human Fecal Sample 5 (H5) (**Supplemental Table 7)**. As mentioned above, these five samples were the only ones to show Bifidobacteriaceae *gyrB* amplification as well (**Supplemental Table 5)**.

Additionally, the ZymoBIOMICS Gut Microbiome Standard was used to evaluate off-target amplifications (**Supplemental Table 8**). The Moeller short-read *gyrB* primer sets for Bifidobacteriaceae and Lachnospiraceae generated ASVs not present in the 16S results for the ZymoBIOMICS Gut Microbiome Standard (**Supplemental Table 8**). The Nichols long-read Bacteroidaceae *gyrB* primer set had zero off-target amplifications. Both the Nichols long-read *gyrB* primer sets for Bifidobacteriaceae (2), and Lachnospiraceae (3) had several off-target amplifications confirmed to be present the 16S results for the ZymoBIOMICS Gut Microbiome Standard (**Supplemental Table 8**). None of the Nichols long-read *gyrB* primer sets had additional off-target amplifications (**Supplemental Table 8**).

## Discussion

The use of short-read *gyrB* primers were successfully used in the past to investigate cospeciation between the bacterial members of the gut microbiome and human hosts [11]. In the meantime, there have been technological improvements in sequencing that can be used to further elucidate these relationships. Here, we generated and validated primers designed for long-read *gyrB* sequencing targeting the Bacteroidaceae, Bifidobacteriaceae, and Lachnospiraceae families via PacBio sequencing and compared them to the existing short-read *gyrB* primer sets.

Long-read sequencing with PacBio utilizes circular consensus sequencing (CCS), which involves sequencing the amplicon multiple times and then taking a consensus to filter out any errors. The PacBio error rate is dependent on how many times the amplicon is sequenced. Since genes like 16S and *gyrB* are relatively small (∼1550 and ∼2500 base pairs respectively), PacBio can sequence these amplicons 8-10 times via CCS, which equates to an error rate of 0.001% [26]. This is superior to the typical error rates in short-read sequencers, which range from ∼0.1 – 0.5 [27]. So not only does PacBio sequencing provide more genetic information of the gene of interest through increased read lengths, but it also provides a lower error rate than short-read technology. This is beneficial, as single nucleotide resolution is necessary for evolutionary and co-evolutionary studies.

Using long-read sequencing technology by design provides more genetic information because the entire gene of interest can be sequenced rather than just a part of it. The longer reads also provide a more accurate analysis of the gene of interest. The current study illustrated this by comparing the Nichols long-read primers with the Moeller short-read primer sets. Both *gyrB* primer sets were used to sequence the same eight samples (five human samples [H1-H5], two murine samples [M1 and M2] and a mock community sample of known composition [GM]). It should be noted that both the Nichols long-read and the Moeller short-read primers resulted in some off target amplification, but the rate of off-target amplification was lower for the Nichols long-read primer sets (0%, 0.89% and 10.43% compared to 1.09%, 13.68% and 23.24% for the Bacteroidaceae, Bifidobacteriaceae, and Lachnospiraceae families respectively). Additionally, the Nichols long-read primers showed a higher specificity of the ASVs when compared to the pseudo-MiSeq sequences. On a sample-by-sample basis, the Nichols long-read primers produced a higher percentage of on-target reads when compared to the Moeller short-read primer sets (with the exception of the Lachnospiraceae in the mock sample, which did perform better with the Moeller short-read primer sets).

Overall, our newly created long-read *gyrB* primer sets and PacBio library preparation protocol provide a more robust sequencing approach for this non-standard phylogenetic marker gene compared to published short-read sequencing. This improved accuracy and confidence in the classification will enable greater resolution and specificity for cospeciation studies moving forward.

## Data availability

All sequence data is available on SRA under Bioproject ID PRJNA1055511. All code is available on GitHub at https://github.com/davenport-lab/Long-read_gyrB_method. Reference databases used for the analyses are available on Zenodo under DOIs: 10.5281/zenodo.10451935, 10.5281/zenodo.10452184, and 10.5281/zenodo.10452279. A detailed protocol for generating PacBio libraries is available at protocols.io at dx.doi.org/10.17504/protocols.io.36wgq31nylk5/v1.

## Acknowledgements

The co-authors would like to acknowledge the Huck Institutes’ Genomics Core Facility (RRID:SCR_023645) for sequencing data generation.

---

**Supplemental Figure 1: Tanglegram of the *gyrB* maximum likelihood trees created from the Bifidobacteriaceae PacBio ASVs and the Bifidobacteriaceae pseudo-MiSeq ASVs.** Bifidobacteriaceae phylogeny was generated using the maximum likelihood (ML) method on ASV data collected across all samples. ML trees for long-read PacBio ASVs (left) and short-read pseudo-MiSeq ASVs (right) were compared with a tanglegram. Each tip is annotated with the sample it originated from, the ASV number, and the taxonomic information after classification. ASV classification names are colored by species. The black lines connecting the two trees illustrate a 1-to-1 relationship where the pseudo-MiSeq ASV results in the same classification as the PacBio parent read. The red dashed lines illustrate a scenario where the pseudo-MiSeq sequence has a less specific classification than the parent PacBio sequence. The blue dashed line represents conflicting classifications between the pseudo-MiSeq ASV and its parent PacBio ASV. The normalized Robinson-Fould distance for these two Bifidobacteriaceae trees was 0.33.

**Supplemental Figure 2: Tanglegram of the *gyrB* maximum likelihood trees created from the Lachnospiraceae PacBio ASVs and the Lachnospiraceae pseudo-MiSeq ASVs.** Lachnospiraceae phylogeny was generated using the maximum likelihood (ML) method on ASV data collected across all samples. ML trees for long-read PacBio ASVs (left) and short-read pseudo-MiSeq ASVs (right) were compared with a tanglegram. Each tip is annotated with the sample it originated from, the ASV number, and the taxonomic information after classification. ASV classification names are colored by species. The black lines connecting the two trees illustrate a 1-to-1 relationship where the pseudo-MiSeq ASV results in the same classification as the PacBio parent read. The red dashed lines illustrate a scenario where the pseudo-MiSeq sequence has a less specific classification than the parent PacBio sequence. The blue dashed line represents conflicting classifications between the pseudo-MiSeq ASV and its parent PacBio ASV. The normalized Robinson-Fould distance for these two Lachnospiraceae trees was 0.40.

**Supplemental Figure 3: The *gyrB* maximum likelihood tree for Bacteroidaceae created from short-read MiSeq ASVs.** 82 *gyrB* ASVs were classified from the Bacteroidaceae MiSeq reads. Tips of the tree are ASVs from each sample, labeled first with the sample identifier they came from, the ASV label, and the taxonomic classification of that ASV. ASVs from the same species are colored identically for visualization with off-target ASVs colored in black. Of the 82 *gyrB* ASVs, 55 ASVs classified to Bacteroidaceae.

**Supplemental Figure 4: The *gyrB* maximum likelihood tree for Bacteroidaceae created from long-read PacBio ASVs.** 31 *gyrB* ASVs were classified from the Bacteroidaceae PacBio reads. Tips of the tree are ASVs from each sample, labeled first with the sample identifier they came from, the ASV label, and the taxonomic classification of that ASV. ASVs from the same species are colored identically for visualization. All 31 *gyrB* ASVs classified to Bacteroidaceae.

**Supplemental Figure 5: The *gyrB* maximum likelihood tree for Bifidobacteriaceae created from long-read PacBio ASVs.** 66 *gyrB* ASVs were classified from the Bifidobacteriaceae PacBio reads. Tips of the tree are ASVs from each sample, labeled first with the sample identifier they came from, the ASV label, and the taxonomic classification of that ASV. ASVs from the same species are colored identically for visualization. The complete ML tree containing all 66 ASVs can be seen on the left. Of the 66 *gyrB* ASVs, only 26 ASVs classified to Bifidobacteriaceae. Off target ASVs will remain black

**Supplemental Figure 6: The *gyrB* maximum likelihood tree for Lachnospiraceae created from the short-read MiSeq ASVs.** 367 *gyrB* ASVs were classified from the Lachnospiraceae MiSeq reads. Tips of the tree are ASVs from each sample, labeled first with the sample identifier they came from, the ASV label, and the taxonomic classification of that ASV. ASVs from the same species are colored identically for visualization, with off-target ASVs colored in black. Of the 367 *gyrB* ASVs, 250 ASVs classified to Lachnospiraceae.

**Supplemental Figure 7: The *gyrB* maximum likelihood tree for Lachnospiraceae created from long-read PacBio ASVs.** 456 *gyrB* ASVs were classified from the Lachnospiraceae PacBio reads. Tips of the tree are ASVs from each sample, labeled first with the sample identifier they came from, the ASV label, and the taxonomic classification of that ASV. ASVs from the same species are colored identically for visualization, with off-target ASVs colored in black. Of the 456 *gyrB* ASVs, 326 ASVs classified to Lachnospiraceae.

**Supplemental Table 1: Candidate Bacteroidaceae long-read amplicon primers generated by PhyloTAGs.** PhyloTAGs generated a total of 1,953 potential primer locations using a 21bp sliding window approach across the Bacteroidaceae *gyrB* gene, including forward and reverse primers for each location. Position_window_(bp) indicates the location of the sliding window, Degeneracies lists the total number of degeneracies within that window, Bases_per_position lists the observed bases per position in the reference database, Forward_primer is the proposed forward primer that matches that location, and Reverse_primer is the proposed reverse primer that matches that location.

**Supplemental Table 2: Candidate Bifidobacteriaceae long-read amplicon primers generated by PhyloTAGs.** PhyloTAGs generated a total of 2,121 potential primer locations using a 21bp sliding window approach across the Bifidobacteriaceae *gyrB* gene, including forward and reverse primers for each location. Position_window_(bp) indicates the location of the sliding window, Degeneracies lists the total number of degeneracies within that window, Bases_per_position lists the observed bases per position in the reference database, Forward_primer is the proposed forward primer that matches that location, and Reverse_primer is the proposed reverse primer that matches that location.

**Supplemental Table 2: Candidate Lachnospiraceae long-read amplicon primers generated by PhyloTAGs.** PhyloTAGs generated a total of 1,932 potential primer locations using a 21bp sliding window approach across the Lachnospiraceae *gyrB* gene, including forward and reverse primers for each location. Position_window_(bp) indicates the location of the sliding window, Degeneracies lists the total number of degeneracies within that window, Bases_per_position lists the observed bases per position in the reference database, Forward_primer is the proposed forward primer that matches that location, and Reverse_primer is the proposed reverse primer that matches that location.

**Supplemental Table 4:** Breakdown of the sequenced reads for both primer sets for the Bacteroidaceae family: An in-depth break down of the on- and off- target *gyrB* reads for each sample and primer set for the Bacteroidaceae amplicons. “On target” is defined as reads or ASVs classifying to the bacterial family targeted by the primer, while “off target” is defined as reads or ASVs classifying to any other bacterial family.

|  | GM | H1 | H2 | H3 | H4 | H5 | M1 | M2 | Total | ASV |
| --- | --- | --- | --- | --- | --- | --- | --- | --- | --- | --- |
| On target Reads (Moeller Primers) | 94983 | 129383 | 97876 | 67539 | 56478 | 120293 | 112839 | 124722 | 804113 | 55 |
| Off-target Reads (Moeller Primers) | 24 | 31 | 187 | 302 | 1188 | 7080 | 21 | 39 | 8872 | 27 |
| Total number of reads (Moeller Primers) | 95007 | 129414 | 98063 | 67841 | 57666 | 127373 | 112860 | 124761 | 812985 | N/A |
| Percent of reads on target (Moeller Primers) | 99.97% | 99.98% | 99.81% | 99.55% | 97.94% | 94.44% | 99.98% | 99.97% | 98.91% | N/A |
| On target Reads (Nichols Primers) | 37767 | 50369 | 38751 | 52891 | 52644 | 22274 | 46910 | 44941 | 346547 | 31 |
| Off-target Reads (Nichols Primers) | 0 | 0 | 0 | 0 | 0 | 0 | 0 | 0 | 0 | 0 |
| Total number of reads (Nichols Primers) | 37767 | 50369 | 38751 | 52891 | 52644 | 22274 | 46910 | 44941 | 346547 | N/A |
| Percent of reads on target (Nichols Primers) | 100% | 100% | 100% | 100% | 100% | 100% | 100% | 100% | 100% | N/A |

**Supplemental Table 5:** Breakdown of the sequenced reads for both primer sets for the Bifidobacteriaceae family: An in-depth break down of the on- and off-target *gyrB* reads for each sample and primer set for the Bifidobacteriaceae amplicons. “On target” is defined as reads or ASVs classifying to the bacterial family targeted by the primer, while “off target” is defined as reads or ASVs classifying to any other bacterial family. Only samples that showed amplification after gel electrophoresis were included in this analysis.

|  | GM | H1 | H2 | H4 | H5 | Total | ASVs |
| --- | --- | --- | --- | --- | --- | --- | --- |
| On target Reads (Moeller Primers) | 141892 | 171261 | 72252 | 58687 | 166518 | 610610 | 28 |
| Off target Reads (Moeller Primers) | 1166 | 12066 | 56230 | 18892 | 8437 | 96791 | 228 |
| Total number of reads (Moeller Primers) | 143058 | 183327 | 128482 | 77579 | 174955 | 707401 | N/A |
| Percent of reads on target (Moeller Primers) | 99.18% | 93.42% | 56.24% | 75.65% | 95.18% | 86.32% | N/A |
| On target Reads (Nichols Primers) | 52382 | 40621 | 41513 | 101523 | 23715 | 259754 | 26 |
| Off target Reads (Nichols Primers) | 141 | 868 | 978 | 23 | 332 | 2342 | 40 |
| Total number of reads (Nichols Primers) | 52523 | 41489 | 42491 | 101546 | 24047 | 262096 | N/A |
| Percent of reads on target (Nichols Primers) | 99.73% | 97.91% | 97.70% | 99.98% | 98.62% | 99.10% | N/A |

**Supplemental Table 6:** Breakdown of the sequenced reads for both primer sets for the Lachnospiraceae family: An in-depth break down of the on- and off- target *gyrB* reads for each sample and primer set for the Lachnospiraceae amplicons. “On target” is defined as reads or ASVs classifying to the bacterial family targeted by the primer, while “off target” is defined as reads or ASVs classifying to any other bacterial family

|  | GM | H1 | H2 | H3 | H4 | H5 | M1 | M2 | Total | ASV |
| --- | --- | --- | --- | --- | --- | --- | --- | --- | --- | --- |
| On target Reads (Moeller Primers) | 213816 | 43997 | 114592 | 105904 | 108617 | 109306 | 107495 | 94674 | 898401 | 250 |
| Off-target Reads (Moeller Primers) | 309 | 83804 | 48463 | 3554 | 9420 | 17012 | 46438 | 62996 | 271996 | 117 |
| Total number of reads (Moeller Primers) | 214125 | 127801 | 163055 | 109458 | 118037 | 126318 | 153933 | 157670 | 1170397 | N/A |
| Percent of reads on target (Moeller Primers) | 99.86% | 34.43% | 70.28% | 96.75% | 92.02% | 86.53% | 69.83% | 60.05% | 76.76% | N/A |
| On target Reads (Nichols Primers) | 36664 | 98197 | 70767 | 57172 | 95671 | 100231 | 96069 | 65942 | 620713 | 326 |
| Off-target Reads (Nichols Primers) | 28589 | 1859 | 5169 | 1920 | 24657 | 4146 | 2602 | 3354 | 72296 | 130 |
| Total number of reads (Nichols Primers) | 65253 | 100056 | 75936 | 59092 | 120328 | 104377 | 98671 | 69296 | 693009 | N/A |
| Percent of reads on target (Nichols Primers) | 56.19% | 98.14% | 93.19% | 96.75% | 79.51% | 96.03% | 97.36% | 95.16% | 89.57% | N/A |

**Supplemental Table 7:** 16S rRNA gene sequencing confirms the presence or absence of Bacteroidaceae, Bifidobacteriaceae, and Lachnospiraceae. The 16S rRNA gene sequencing results are broken down to illustrate how many reads were present for the Bacteroidaceae, Bifidobacteriaceae, and Lachnospiraceae bacterial families in each sample. Total reads are provided for each sample as well.

|  | GM | H1 | H2 | H3 | H4 | H5 | M1 | M2 |
| --- | --- | --- | --- | --- | --- | --- | --- | --- |
| Total Bacteroidaceae Reads | 4830 | 772 | 1845 | 6905 | 72 | 145 | 198 | 197 |
| Total Bifidobacteriaceae Reads | 13 | 960 | 49 | 0 | 1674 | 3327 | 0 | 0 |
| Total Lachnospiraceae Reads | 320 | 6731 | 5725 | 1613 | 7689 | 7591 | 7305 | 6937 |
| Total 16S reads | 36909 | 18288 | 14489 | 22574 | 29965 | 26042 | 20236 | 20579 |

**Supplemental Table 8:** 16S rRNA gene sequencing of the ZymoBIOMICS Gut Microbiome Standard (GM) confirms off-target amplification of *gyrB* primers. The first column (“Taxa”) lists all the bacterial genera identified with 16S sequencing of the ZymoBIOMICS Gut Microbiome Standard (“16S”). The following columns list whether the listed genus was identified with the specific primer sets indicated. The ‘Unclassified’ row is used to mark the presence or absence of unclassified genera. The ‘Other’ row is used to mark the presence or absence of genera that were not observed in the 16S rRNA gene analysis of the ZymoBIOMICS Gut Microbiome Standard but were identified by the *gyrB* amplicon analysis. “Off-target” amplification is marked with asterisks***.

| Taxa | 16S | Bacteroidaceae<br>Moeller MiSeq<br>*gyrB* | Bacteroidaceae<br>Nichols PacBio<br>*gyrB* | Bifidobacteriaceae<br>Moeller MiSeq *gyrB* | Bifidobacteriaceae<br>Nichols PacBio *gyrB* | Lachnospiraceae<br>Moeller MiSeq *gyrB* | Lachnospiraceae Nichols<br>PacBio *gyrB* |
| --- | --- | --- | --- | --- | --- | --- | --- |
| Akkermansia | Present | NotPresent | NotPresent | NotPresent | NotPresent | NotPresent | NotPresent |
| Asaccharospora | Present | NotPresent | NotPresent | NotPresent | NotPresent | NotPresent | NotPresent |
| Bacteroides | Present | Present | Present | Present*** | NotPresent | NotPresent | Present*** |
| Bifidobacterium | Present | NotPresent | NotPresent | Present | Present | NotPresent | Present*** |
| Escherichia/<br>Shigella | Present | NotPresent | NotPresent | NotPresent | Present*** | NotPresent | NotPresent |
| Faecalibacterium | Present | NotPresent | NotPresent | Present*** | Present*** | NotPresent | Present*** |
| Fusobacterium | Present | NotPresent | NotPresent | NotPresent | NotPresent | NotPresent | NotPresent |
| Lactobacillus | Present | NotPresent | NotPresent | NotPresent | NotPresent | NotPresent | NotPresent |
| Leminorella | Present | NotPresent | NotPresent | NotPresent | NotPresent | NotPresent | NotPresent |
| Peptoclostridium | Present | NotPresent | NotPresent | NotPresent | NotPresent | Present*** | NotPresent |
| Prevotella | Present | NotPresent | NotPresent | Present*** | NotPresent | NotPresent | NotPresent |
| Roseburia | Present | NotPresent | NotPresent | Present*** | NotPresent | Present | Present |
| Veillonella | Present | NotPresent | NotPresent | NotPresent | NotPresent | NotPresent | NotPresent |
| Unclassified | Not<br>Present | Present*** | NotPresent | NotPresent | NotPresent | Present*** | NotPresent |
| Other | Not<br>Present | NotPresent | NotPresent | Present*** | NotPresent | Present*** | NotPresent |

